# Plume dynamics structure the spatiotemporal activity of glomerular networks in the mouse olfactory bulb

**DOI:** 10.1101/2020.11.25.399089

**Authors:** Suzanne M. Lewis, Lai Xu, Nicola Rigolli, Mohammad F. Tariq, Merav Stern, Agnese Seminara, David H. Gire

## Abstract

Although mice locate resources using turbulent airborne odor plumes, the stochasticity and intermittency of fluctuating plumes create challenges for interpreting odor cues in natural environments. Population activity within the olfactory bulb (OB), is thought to process this complex spatial and temporal information, but how plume dynamics impact odor representation in this early stage of the mouse olfactory system is not known. Limitations in odor detection technology have made it impossible to measure plume fluctuations while simultaneously recording from the mouse’s brain. Thus, previous studies have measured OB activity following controlled odor pulses of varying profiles or frequencies, but this approach only captures a subset of features found within olfactory plumes. Adequately sampling this feature space is difficult given a lack of knowledge regarding which features the brain extracts during exposure to natural olfactory scenes. Here we measured OB responses to naturally fluctuating odor plumes using a miniature, adapted odor sensor combined with wide-field GCaMP6f signaling from the dendrites of mitral and tufted (MT) cells imaged in olfactory glomeruli of head-fixed mice. We precisely tracked plume dynamics and imaged glomerular responses to this fluctuating input, while varying flow conditions across a range of ethologically-relevant values. We found that a consistent portion of MT activity in glomeruli follows odor concentration dynamics, and the strongest responding glomeruli are the best at following fluctuations within odor plumes. Further, the reliability and average response magnitude of glomerular populations of MT cells are affected by the flow condition in which the animal samples the plume, with the fidelity of plume following by MT cells increasing in conditions of higher flow velocity where odor dynamics result in intermittent whiffs of stronger concentration. Thus, the flow environment in which an animal encounters an odor has a large-scale impact on the temporal representation of an odor plume in the OB. Additionally, across flow conditions odor dynamics are a major driver of activity in many glomerular networks. Taken together, these data demonstrate that plume dynamics structure olfactory representations in the first stage of odor processing in the mouse olfactory system.

## Introduction

Mice are adept at localizing odor sources ((Gire et al., 2016); (Baker et al., 2018); (Liu et al., 2019); (Gumaste et al., 2020)), but the spatiotemporal information in olfactory environments that aids this search behavior is unknown. Odors travel in plumes which pull odor away from its source in filaments that are broken and distorted as they travel in air, creating complex odor environments. From the perspective of an olfactory searcher, these intermittent filaments create stochastic odor encounters, or whiffs, such that odor concentration dynamics fluctuate rapidly from moment to moment. Features of these complex plume dynamics contain information regarding odor source location ((Murlis et al., 2000); (Celani et al., 2014)). For example, as a searcher encounters odors, the frequency, strength, and timing of encounters provide complex cues about an odor source ((Vergassola et al., 2007); (Ache et al., 2016); (Michaelis et al., 2020); (Atema, 1996). A simple strategy such as averaging odor concentration across whiffs could eliminate the complexity of an odor plume, allowing an animal to simply follow an increasing odor concentration gradient to the odor source. However, an animal dependent on this search strategy would operate at a timescale far slower than that observed in mice engaged in olfactory-guided search (Gumaste et al., 2020). This suggests that rodents most likely extract information from the complex spatiotemporal dynamics of olfactory environments to support their efficient odor-guided search behavior.

The extraction of information from fluctuating odor plumes will necessarily be impacted by the physics of odor transport. Factors such as wind speed and the Reynolds number of the plume could impact early olfactory processing in mammals. Precise olfactometers have been used to model certain features found in natural odor environments such as fluctuating and intermittent odor concentration dynamics. Although this work provides important insights, olfactometers do not capture the full complexity of the odor environment. One problem is a lack of knowledge regarding which features of the plume are relevant to olfactory search, constraining which features olfactometers have been used to mimic. In addition, olfactometers create artificial plumes that decouple odor concentration from features present in olfactory environments. This omits correlations between concentration fluctuations and the surrounding air flow as well as small scale details of odor transport like diffusion. Decoupling these factors creates challenges for interpretation because it implicitly disrupts processing moderated by these features, such as the impact of wind speed on the vibrissal system ((Yu et al., 2016)) or feedback regarding bilateral nasal sampling ((Esquivelzeta Rabell et al., 2017); (Markopoulos et al., 2012)). Directly observing how MT activity is impacted by the plume dynamics of natural olfactory scenes will thus constrain hypotheses regarding which spatiotemporal features of natural odor stimuli are conveyed by the brain.

We studied the response of MT cells in the OB to odor concentration dynamics in awake mice as they processed natural olfactory scenes, i.e. odor plumes. We used wide-field calcium imaging to measure MT activity, allowing us to study OB output at the level of glomerular complexes on the dorsal surface of the OB. Simultaneous recordings of the OB and plume dynamics show glomerular population activity follows fluctuations of odor concentration during plume encounters. The reliability and following behavior of glomerular responses were moderated by wind speed and the resulting changes in plume structure. The fidelity of odor concentration tracking increased when concentration dynamics were skewed, creating intermittent odor encounters across the plume presentation. In addition, as the strength and reliability of odor-evoked activity in MT cells increased, this activity more accurately followed plume dynamics. Together, these data demonstrate for the first time that the rapid fluctuations present in natural olfactory scenes significantly structure the activity of glomerular MT cell populations in the mouse OB.

## Materials and Methods

### Olfactory Stimuli

Olfactory stimuli were released by an automated odor port within a 32” X 16” X 16” acrylic wind tunnel where airspeed was controlled by a vacuum at the rear of the wind tunnel, posterior to the animal’s location. Concentration dynamics of olfactory stimuli varied stochastically from trial to trial creating plumes with unique concentration dynamics on each trial. Adjusting the velocity of wind flow allowed for variation in the Reynolds number, resulting in characteristic changes of plume dynamics for different flow levels (Supplemental Figure 1). Reynolds numbers were calculated using the mean flow of each condition and the half height of the tunnel (8”) as the head-fix setup is placed on a stage that is elevated approximately 20cm from tunnel floor. Low, medium, and high flow Reynolds numbers were 2400 *±* 1000, 8800 *±* 400, and 9800 *±* 200 respectively (mean *±* st dev). Absolute velocities of low, medium and high flow conditions were .40 *±* 0.16, 1.31 *±* 0.05, and 1.81 *±* 0.17 and velocity fluctuations were 14 *±* 3 fpm, 6 *±* 2 fpm, and 5.3 *±*1.5 fpm respectively.

A single session consisted of forty trials of odor presentation. Odor ports were located upwind *~* 13 cm anterior to the animal’s nose, and each plume presentation had a duration of 10 seconds. Odor release began approximately 10 seconds into the trial. In order to avoid responses predicting the beginning of the plume, the exact time of odor release was jittered by adjusting the duration of the 5 second intertrial interval by a length of time drawn randomly from a uniform distribution, *U* (*−* 2, 2). Random clicking noise was used to control for the clicking sound of the port serving as a cue for plume onset. Starting 5 seconds prior to plume onset, a number was drawn from a uniform distribution, *U* (0, 1), for each camera frame, and if the number exceeded 0.95 a clicking sound was produced. The odor used was a mixture of benzaldehyde and ethanol. Ethanol concentration throughout each trial was measured by a modified, commercially available ethanol sensor placed within 3.5-4mm from the mouse’s right nostril.

### Implantation of Cranial Window

Implantation of the imaging window was adapted from methodology detailed in Batista-Brito et al., (Batista-Brito et al., 2017). Mice were anesthetized with isoflurane for surgery. 2 × 2.5mm or 2 × 3mm craniotomies were performed above the olfactory bulbs and custom cut double windows were implanted. A customized stainless steel head plate was glued directly on the skull posterior to the window, and two Invivo1 stainless steel screws (McMaster-Carr) were placed posterior to the head plate. Metabond was then added to cover all exposed skull and a thin layer built to cover the screws and the central surface of the headplate. The position of the craniotomy was biased towards either the left or right bulb.

### In vivo imaging

Widefield fluorescent microscopy was used for awake, head-fixed imaging in Thy1-GCaMP6f-GP 5.11 (IMSR Cat JAX:024339, RRID:IMSR JAX:024339) mice to view neural activity in the dorsal OB. Mice were between 11 weeks-13 months old when imaged. 488 nm LED stimulation was used for the duration of the trial (30s), but was absent during intertrial intervals (*~* 5 seconds) to avoid excessive bleaching. All mice were imaged at 30Hz with 4x 0.13 NA objective (Nikon). Neural activity was recorded using a Teledyne Photometrics Prime 95B sCMOS camera. For each session, mice were head-fixed above a freely rotating, circular track, allowing mice to run at will during imaging sessions.

### In vitro OB slice imaging

To establish patterns of expression and signals obtained from the OB of Thy1-GCaMP6f-GP 5.11 animals, imaging experiments were conducted in OB slices. Horizontal OB slices (300–400 m) were made following isoflurane anesthesia and decapitation. Olfactory bulbs were rapidly removed and placed in oxygenated (95% O2, 5% CO2) ice-cold solution containing the following (in mM): 83 NaCl, 2.5 KCl, 3.3 MgSO4, 1 NaH2PO4, 26.2 NaHCO3, 22 glucose, 72 sucrose, and 0.5 CaCl2. Olfactory bulbs were separated into hemispheres with a razor blade and attached to a stage using adhesive glue applied to the ventral surface of the tissue. Slices were cut using a vibrating microslicer (Leica VT1000S) and were incubated in a holding chamber for 30 min at 32°C. Subsequently, the slices were stored at room temperature. Slices were placed on a Scientifica SliceScope Pro 6000, using near infrared imaging for slice placement and 488 nm LED illumination for imaging activity and a QI825 Scientific CCD Camera (Q Imaging) for image acquisition. Imaging was performed at 32–35°C. The base extracellular solution contained the following: 125 mM NaCl, 25 mM NaHCO3, 1.25 mM NaHPO4, 25 mM glucose, 3 mM KCl, 1 mM MgCl2, and 2 mM CaCl2 (pH 7.3 and adjusted to 295 mOsm), and was oxygenated (95% O2, 5% CO2). An elevated KCl solution (equimolar replacement of 50 mM NaCl with KCl in the extracellular solution) locally applied through a borosilicate pipette using a picospritzer 2 (Parker Instrumentation) was used to stimulate cells.

### Data Preprocessing

ImageJ was used to crop fields of view (FOVs) for data analysis. It was also used to extract pixel averaged signal for hand-drawn region of interest (ROI) anaylsis.

Matlab 2019b was used to analyze data and plot figures. Data was aligned using NoRMCorre software to perform piecewise rigid and non-rigid motion correction (Pnevmatikakis and Giovannucci (2017)). This alignment corrected for both global frame movement due to head jitter (rigid) and localized distortion due to brain movement (non-rigid).

### Precise tracking of plume dynamics

A miniaturized ethanol sensor modified from the commercially available metal oxide (MOX) sensors (Tariq et al., 2019) was placed within 4mm of the lateral edge of the mouse’s right nostril to capture the odor concentration signal across plume presentations for each trial. A single odor, a benzeldahyde-ethanol mixture, was used for each trial. Odors released together travel together within plumes at sufficiently small scales because dispersion dominates over diffusion ((Celani et al., 2014); (Yeung and Pope, 1993)). Therefore, the ethanol sensor measured the odor concentration of the benzeldahyde-ethanol mixture. The plume for each trial was released by an automated odor port at the upwind end of the wind tunnel. The odor was released 10 seconds after the trial began for a duration of 10 seconds. Each indicated flow condition was maintained throughout the entire trial. Flow condition was set for each trial block (figure 1C) by adjusting the strength of a vacuum exhaust at the downwind side of the wind tunnel to one of three levels, low, medium, and high. A medium level flow was presented for the first 10 trials, after which flow alternated by high and low flow in blocks of 5 trials. Some sessions transitioned from medium to low flow initially and others transitioned from medium to high flow.

**Figure 1.**
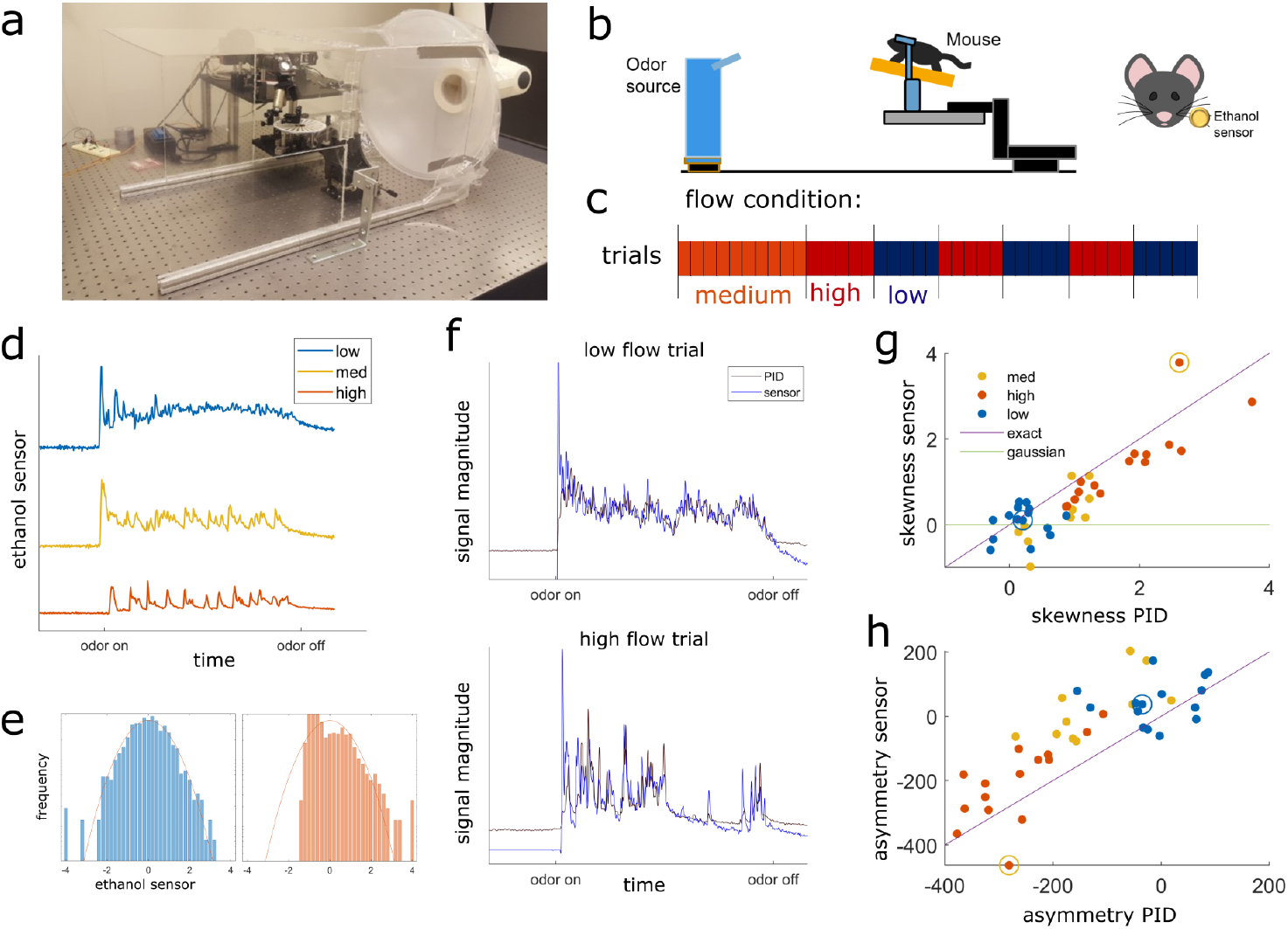
Plume presentations and head-fix setup for in-vivo recording experiments. a) All experiments conducted in 16” × 16” × 32” wind tunnel for quick clearing of odor presentations. The odor port (not pictured) was located ~ 13 cms upwind of the animal’s nose. b) (right) Graphic detailing experimental setup. (left) Ethanol odor concentration measured using a modified, commercially available ethanol sensor placed ~ 4mm from the edge of the mouse’s right nostril. c) Diagram depicting flow conditions (high, medium, or low) of the 40 trials within a single session. d) Example odor traces are depicted for each flow condition. e) Histograms of the odor concentration magnitude sampled across two examples trials show a change in skewness between low flow (right, blue) and high flow (left, red), with skewness increasing with increased airflow during plume presentations. f) Comparisons between the deconvolved sensor signal and a PID signal during a set of paired recordings show odor concentration dynamics of the deconvolution can recover dynamics observed in the PID recordings (*r* = 0.61, *p* < .001). An example from low flow (top) and high flow (bottom) are shown. g-h) For each trail, the skewness (F) and asymmetry (G) of the plume’s distribution are calculated showing that high flow trials (orange) are separable from low (blue) for both PID and sensor recordings.

### Ethanol Deconvolution

Recent characterization of MOX sesnsors comparing their deconvolved signals to simultaneously recorded photoionization detector (PID) signals have validated the use of MOX sensors in capturing turbulent plume dynamics despite their slower recording dynamics ((Martinez et al., 2019); (Tariq et al., 2019)). For our experiments, ethanol concentration throughout each trial was measured by a modified, commercially available ethanol sensor placed within 3.5-4mm from the mouse’s right nostril. A single session consisted of 40 trials of odor presentation. Odor ports were located upwind *~* 13 cm anterior to the animal’s nose, and each presentation had a duration of 10 seconds and consisted of a mixture of benzaldehyde and ethanol.

Sensor signal was acquired at 100Hz and then low pass filtered at 30 Hz using a Kaiser window. The signal, *e*, was then normalized within each trial using the mean and standard deviation of the signal during the plume presentation. The signal was deconvolved by adapting the deconvolution specified in Tariq et al., 2019. The kernel was defined in the same manner, but instead of normalizing the range of the kernel, the integral of the kernel is normalized. Thus, the kernel, *k*, is calculated as follows.

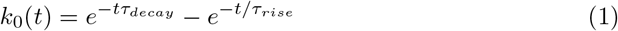

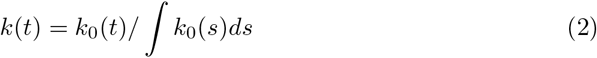

where *t* is an array with evenly spaced timestamps at the proper sampling rate for the length of a single trial, *τ*_*decay*_ = 0.0001, and *τ*_*rise*_ = 0.4629. Both signals, *e* and *k*, are then transformed into Fourier space using the Matlab Fourier transform function, and the ethanol signal is deconvolved by dividing *ê* by 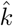. The inverse Fourier transform of the resulting deconvolution is taken to obtain *d*. The deconvolved signal, *d*, is then normalized within each trial using the mean and standard during the plume presentation. The deconvolution optimizes the preservation of odor concentration dynamics across trials, but does not preserve the absolute value of odor concentration.

To optimize the parameters for the deconvolution, a complete session of 40 plume presentations was recorded with the usual ordering of flow condition blocks (figure 1C). No mice were recorded during this session. Instead, a photoionization detector (PID) was placed 4mm from the ethanol sensor in the same position where the animal is usually head-fixed.

To optimize parameters, the PID signal, *p* was first downsampled to 100Hz to match the sensor sampling rate. Next, the signal was normalized within each trial using the mean and standard deviation of the signal during plume presentation. This normalized signal was then compared to *d* deconvolved across a range of *τ*_*decay*_ and *τ*_*rise*_ parameter values. The Kernel parameters were chosen by minimizing mean squared error between the *e* and *p* signals averaged across all trials within the paired recording session. The deconvolved ethanol sensor signal allows for the recovery of plume dynamics unique to each trial (Supplemental Figure 2). It is significantly correlated with the PID signal as measured during plume presentations (*r* = 0.61, *p <* 0.001, both sampled at 100*Hz*), which is a 0.22 improvement from the correlation between the raw ethanol sensor and PID signal (*r* = 0.39, *p <* 0.001, both sampled at 100*Hz*).

Finally, with the exception of Supplemental Figure 1, the deconvolved trace was downsampled for figures and analyses to match the calcium trace (30Hz) by averaging all samples taken across each camera frame.

### Defining dynamic flow

Dynamic flow was calculated within each session. In the experimental setup, flow condition is defined by wind speed. Intermittency was measured within a trial during the middle 8 seconds of the 10 second plume presentation by calculating asymmetry or by using the 3^*rd*^ moment of the sampled distribution. Low and High flow trials were separable using either of these measures.

To determine if there was a main effect of flow condition on these stimulus properties, skewness and asymmetry, a three-way analysis of variance (ANOVAs) test was conducted for each parameter. For each ANOVA, a multiple comparisons using Tukey’s pairwise comparisons was performed to look for significant differences of the parameter between flow where comparisons were considered significant for *p < .*01. These tests indicated that both skewness and asymmetry varies significantly across flow condition (*F* (157) = 52.7, *p < .*001 and *F* (157) = 54.9, *p < .*001 respectively). Multiple comparison tests examining the differential effect of flow on these parameters showed that, for both, there was a significant difference between low and medium flow and a significant difference between low and high flow. No significant difference was found between medium and high flow. Therefore, only low and high flow conditions were selected when examining the effect of air flow on neural parameters.

### Measuring glomerular responses to plume dynamics

Widefield imaging of the dorsal surface of the OB in Thy1-GCaMP6f-GP5.11 mice was used to capture MT cell activity at the glomerular level (figure 2). Thy1 mice exhibit fast kinetics and strong expression in MT cells within the OB (Dana et al., 2014). Global MT activity is clustered into dendritic complexes known as glomeruli. Widespread activity of secondary dendrites in the EPL of dorsal OB (figure 2A) causes diffuse fluorescence across the imaging field. Therefore, CaImAn, a constrained nonnegative matrix factorization(CNMF) algorithm, was used on each FOV to find regions of interest(ROIs) and their activity traces ((Pnevmatikakis et al., 2016)). Thus the spatial decomposition of CNMF provided the ROI for each glomerulus, and the temporal decomposition provided its corresponding denoised activity trace. The denoising of the CNMF temporal decomposition helps to remove correlated signal between neighboring glomeruli, accounts for calcium drift within recording sessions, and separates glomeruli overlapping in the dorsal-ventral dimension. The change in the distribution of correlation coefficients between ROIs before and after de-noising (Supplemental Figure 3) shows a decorrelation of glomerular signals. A two-sample Kolmogorov-Smirnoff test shows the distributions of correlation coefficients between the pixel averaged ROI traces, 0.87 *±* 0.07 (mean ± st. dev), and the CNMF ROI traces, 0.36 *±* 0.28 (mean ± st. dev), are significantly different (*D* = 0.86*, p <* 0.001).

**Figure 2.**
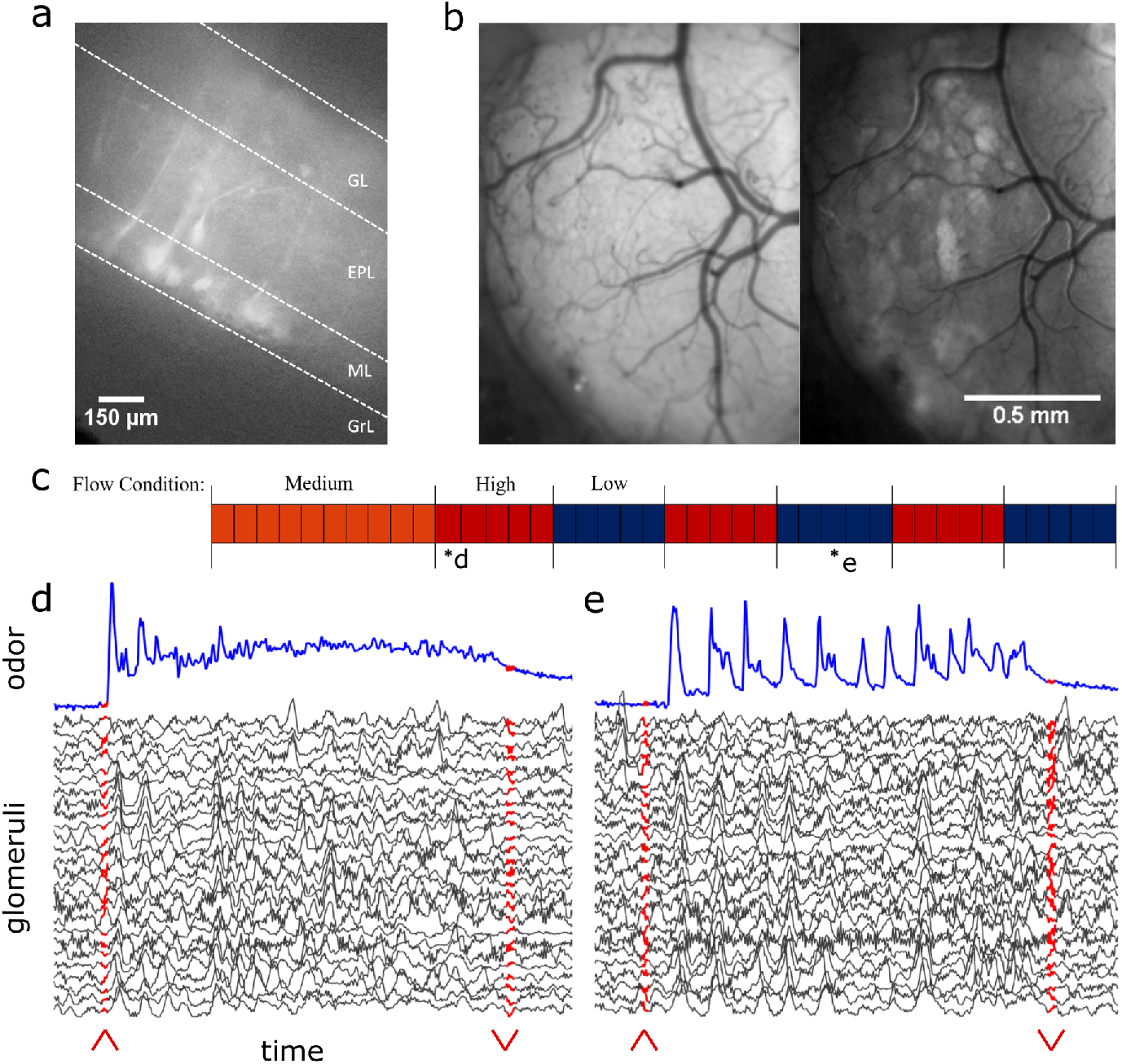
In-vivo recording of glomerular population response. a) In vitro image of MT cell activity in response to K+ puff. b) In vivo view of the dorsal olfactory bulb through an implanted cranial window. (left) Window activity averaged across a single trial. (right) Projected standard deviation for the same trial shows MT activity in the dorsal OB responsive to the odor presentation. c) Diagram depicting flow conditions (high, medium, or low) of the 40 trials within a single session. d) The deconvolved ethanol trace (blue) compared to the deconvolved response of each glomeruli (black) within the recorded FOV during a single low flow trial depicted by asterisk in (c). Red arrows indicate onset and offset of plume presentation. e) Same but for a single high flow trial from the same session also depicted by asterisk in (c).

To protect against oversegmenting a single glomerulus into multilple ROIs, neighboring ROIs whose baseline CNMF activtiy was correlated above 0.65 were selected as candidates for ROI merging. The baseline period was examined as criteria for possible merging since glomeruli might have similar response profiles to the stimulus dynamics during plume presentations. The baseline activity of the neighboring ROIs was then binarized using a threshold of *±*1 st dev. If the correlation of the binarized activity between neighboring ROIs exceeded 0.65, the ROI with the lower mean activity (presumably encompassing less of the glomerulus) was dropped from the analysis.

To validate the use of CNMF at the glomerular level, results were compared to a hand drawn ROI analysis conducted on one of the fields of view (FOV 5) in ImageJ (Supplemental Figure 5). Hand-drawn ROIs were selected after viewing footage and reviewing standard deviation and maximum value projections of activity from the FOV within each trial. The activity averaged from within hand drawn ROIs have higher pairwise correlations than denoised CNMF activity traces. This mirrors what is seen when pixel-averaged activity from within CNMF ROIs (without denoising) is compared to the denoised traces. Thus, the denoising of CNMF increases the spatiotemporal resolution of glomerular activtiy observed in the dorsal OB recordings in both types of analyses. A two-sample Kolmogorov-Smirnoff test shows the distributions of correlation coefficients between the deconvoled hand-drawn ROIs, 0.81 *±* 0.08 (mean *±* st. dev), and the deconvoled CNMF traces, 0.1935 *±* 0.3471 (mean *±* st. dev), are significantly different (*D* = 0.86, *p <* 0.001). Results of the hand-drawn ROI analysis (correlation and power analyses) were qualitatively similar to those found using CNMF corroborating the ability of glomerular networks to resolve odor concentration dynamics. Cross-correlations show a relation between glomerular and ethanol signals during odor presentation with all glomeruli having significant correlation with the plume during odor presentation as compared to their respective null distributions from trial shuffled correlation anaylses. In addition, a strong correlation between glomerular response power (0-5 Hz) and ability to track odor concentration dynamics is also present in the hand-drawn ROIs (*r* = 0.80, *p* = *<* 0.001). Thus, we find that CNMF captures the relationship between glomeruli and plume dynamics while improving the resolution of glomerular network activity and inter-glomerular temporal dynamics.

The CNMF activity traces from the identified glomeruli were baseline normalized using the mean and standard deviation of a 5 second baseline activity period prior to stimulus onset. Traces for each glomerulus were then deconvolved in the style of Stern et al., 2020 to recover the average activity rate of each glomerulus. To find the optimal penalty parameter, *λ*, for deconvolution, *λ* was optimized within each glomerulus. Then the median of this optimized distribution was used as the *λ* in the deconvolution for all glomeruli. After deconvolution, traces were standardized using the standard deviation of the glomerulus’ entire trace. Deconvolved signals were standardized in this way since the florescence range of a glomerulus’s response depends on the number of expressing MT cells, the depth of the glomerulus from the dorsal surface, and other methodological factors unrelated to the magnitude of the response.

### Testing for responsive glomeruli

A glomerulus was considered to be responsive to odor if its deconvolved trace exceeded threshold more time points than expected by chance during plume presentation (figure 7). Since this preliminary measure does not rely on stimulus dynamics, it captures glomeruli that respond to the plume even if their response is unrelated to odor concentration dynamics or only present for part of the plume.

First, the deconvolved trace of a single glomerulus was split into two periods: baseline activity and odor response. The baseline period is a 5 second period at the beginning of each trial prior to plume onset. The odor response period is the time during which the plume is present as well as one second immediately following the plume since inhibition has been shown to induce excitatory rebound responses in tufted cells (Cavarretta et al., 2018). The signal is first baseline normalized by subtracting the mean and dividing by the st dev of the baseline activity within each trial. Next, it is binarized, thresholding for time points where activity exceeded the 95% confidence interval of the glomerulus’s original baseline activity (thresholded at *±*1.96 baseline mean). In this way, each time point that crossed the threshold was considered an event. Within each trial plume presentation, if the number of events exceeded the null expectation (5% of the total number of time points during plume presentation rounded up to the nearest integer), the glomerulus was considered to be responsive to the plume during that trial. The proportion of trials to which the glomerulus responded was calculated within three sets of flow conditions (all flows, low flow, and high flow). The cumulative responsivity score plotted in (figure 7A) is the sum of responsivity scores for all three flow conditions where the contribution of each flow is plotted in different colors as a stacked bar graph.

### Cross-correlation between plume dynamics and corresponding neural responses

To understand the relation between stimulus and response time series, a preliminary analysis was conducted by calculating the correlation coefficients between the two signals for each glomerulus. In the future, more sophisticated techniques will be used to establish how much of the neural representations can be explained by high-fidelity odor concentration encoding.

Due to the stochastic nature of plume onset and offset times, the correlation is only calculated for the middle 8 seconds of the 10 second plume so that onset and offset dynamics are not included and the correlation measure represents the magnitude of plume tracking during plume encounters. The cross-correlation coefficients of a glomerulus *r*_*g*_ is calculated between the ethanol *e* and calcium *c* deconvolutions during plume presentations. For a single glomerulus, the correlation coefficient between the two deconvolutions within a single trial *n* is calculated at all possible lags *l*. Using the xcorr() function in Matlab to compute the coefficients, both signals are mean subtracted prior to calculating the cross-correlations such that the correlation coefficients are synonymous with calculating the pearson correlation coefficient between the two signals at each respective lag value.

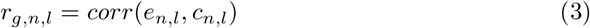

The mean coefficient for each glomerulus, 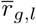 is calculated by averaging across all trials within the session (n=40 for 3 FOVs, and n=39 for 2 FOVs) at each possible lag. The maximum coefficient mean is selected from all lags within a 500ms window *w* of the neural activity following the ethanol signal.

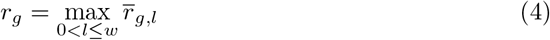

This is considered to be a window of sufficient size to account for variable delays in glomerular processing. The average time lag of *r*_*g*_ was 130ms *±* 100 (mean *±* st. dev).

Within flow cross-correlations are calculated in the same manner but averaged only across trials within the specified flow condition.

For plotting of tracking ability (figure 5A-C), *r*_*g*_ is compared to a single trial shuffled analysis using the same method as detailed above. The difference between the matched and shuffled coefficients suggest correlations are not solely a result of plume structure, but are driven by the temporal dynamics unique to each trial. Trials were shuffled within each glomerulus by calculating the correlations between *e*_*n*_ and *c*_≠*n*_. In this way, any relation dependent on the dynamics of the stochastic fluctuations within each plume presentation is lost, but other statistical features of the plume presentation are preserved, yielding a baseline value for the cross-correlation. Glomeruli are plotted in Figure 5A-C if their correlation coefficient from the matched analysis exceeds *±* 2 standard deviations (st dev of the coefficient distribution from the shuffled analysis) of the shuffled mean coefficient. Since correlation varies significantly within a glomerulus across flow conditions, a glomerulus is considered to exceed the shuffled mean if it does in at least one of the three defined conditions, all, low, or high flow.

Using the same shuffled correlation, a bootstrap analysis was conducted (10,000 iterations) creating a null distribution of the shuffled mean correlation coefficients to test for significance (Supplemental Figure 7c). The mean correlations are compared to their respective 95% confidence interval for the null distribution.

Comparison of correlation coefficients in the matched versus shuffled cross-correlations does not naturally divide the glomeruli into two subpopulations, but rather the strength of this relationship varies continuously across glomeruli. Therefore, instead of dividing glomeruli into subpopulations of tracking versus non-tracking, our analyses consider how the strength of odor concentration tracking compares to other properties of the glomerulus and its response.

## Results

### Measuring glomerular responses to plume dynamics

Using a commercially available, modified odor sensor combined with widefield calcium imaging techniques in head-fixed mice, we reliably tracked plume dynamics and investigated glomerular response to this fluctuating input (figure 1). Imaging was conducted in Thy1-GCaMP6f-GP5.11 mice which have fast kinetics and expression in mitral and tufted (MT) cells within the olfactory bulb (OB) (Dana et al., 2014) (figure 2). Widefield imaging of the dorsal surface of the OB allows for glomerular level resolution of the neural response (figure 2B) (Fletcher et al., 2009). To explore a range of plume dynamics an animal may encounter in its natural environment, we changed the airspeed in the wind tunnel to create stochastic plumes with different odor concentration dynamics (figure 1D-H). Odor identity, concentration and volume released from the odor port remained constant across all flow conditions, making the temporal structure of the plume the only source of variation.

### Mitral and tufted population activity correlates with plume dynamics

At the bulbar level, imaging of MT cell activity shows activation of glomerular networks during odor exposure (figure 3). The global MT activity for each field of view (FOV) was subjected to principal component analysis (PCA) and compared to the simultaneously recorded plume dynamics. The FOV’s were aligned prior to PCA, but no segmentation or denoising was performed. To search for component activity responsive to plume dynamics, the correlation between each principal component and the odor concentration dynamics was calculated. There exists a high ranking component for each mouse that correlates strongly with plume dynamics (figure 3B-D). Plotting the loading weights of the maximally correlated component shows dense clusters of high variance resembling partial spatial maps of glomerular activity. These findings demonstrate that MT population activity recorded in the first relay of olfactory processing is correlated to odor concentration dynamics during plume presentations. In order to establish whether individual glomeruli are correlated to odor cues, we sought to segment the MT activity into glomerular units to determine their respective contributions to the observed tracking of plume fluctuations by population activity.

**Figure 3.**
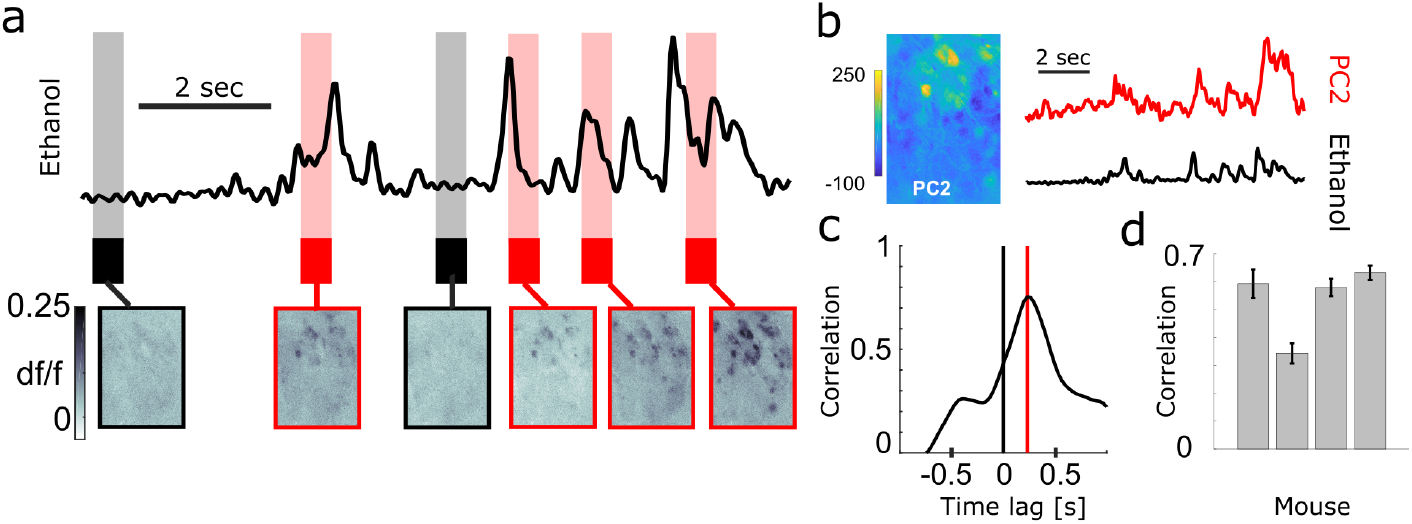
Population response of MT cells in dorsal OB respond to changes in odor concentration during plume presentations. a) Simultaneously recorded deconvolved ethanol plume (top) and imaging of calcium signals from MT cell activity in an example FOV of a Thy1-GCaMP6f (GP5.11) mouse (bottom). Baseline and odorless periods (black) and odor plume input (red) are shown from the indicated time points. Fluctuations in the odor plume elicit repeatable activation of specific glomerular networks in response to whiffs of odor during plume presentations. b) (left) An image of the principal component loadings corresponding to the odor-evoked activity (principal component 2 (PC2)). (right) Time series of PC2 (top, red) aligned to the simultaneous ethanol signal (bottom, black). Scale bar indicates 2 seconds. c) Cross-correlogram between the two signals in (b). Red line indicates a slight offset (~250 ms mean lag across FOVs from sensor to OB response) from 0 for the peak correlation. d) Cross-correlations (mean ± SEM) between odor evoked population activity (principal component) and ethanol sensor signal are strong across 3 Thy1-GCaMP6f (GP5.11) mice (*r* = 0.54 *±* 0.07).

### Neural Activity of Glomeruli

CNMF decomposition provided locations of glomeruli and their corresponding denoised traces (figure 4). To recover the average activity of synaptic complexes of MT acvitity known as glomeruli, CNMF traces were deconvolved in the style of Stern et al., 2020 (Stern et al., 2020)(see methods). The mean deconvolved trace across trials was calculated for each glomerulus during plume presentations. The mean was only considered during the middle 8 seconds of the 10 second odor plume to concentrate on glomerular responses to odor concentration dynamics during plume presentations and avoid responses to onset or offset plume dynamics.

**Figure 4.**
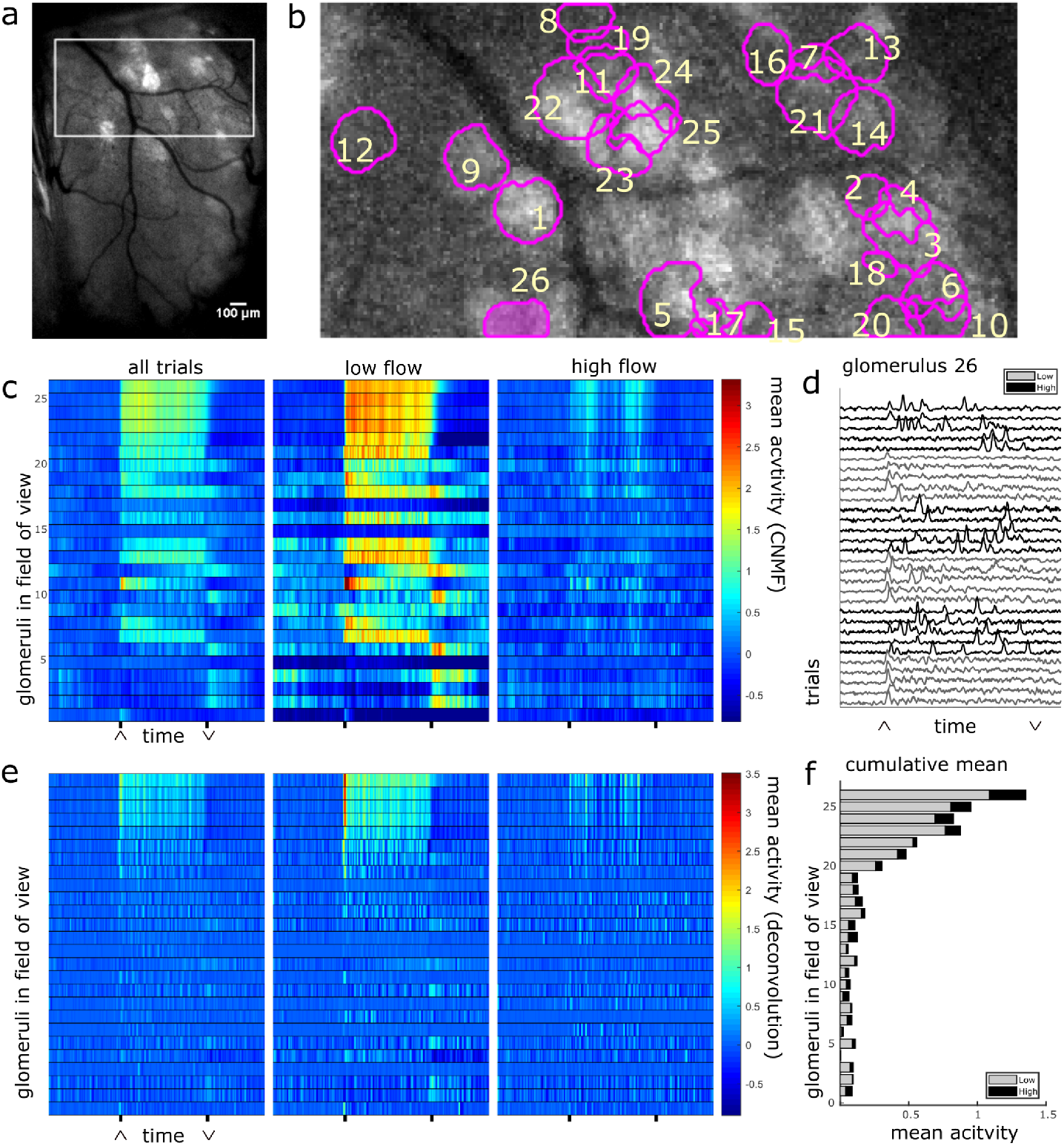
The spatial and temporal decomposition of CNMF identifies glomeruli and denoises their traces. a) The white box outlines the FOV used for analysis as it relates to the larger recording window. The image shows the standard deviation projection of the aligned recording during a single odor presentation. b) Mean subtracted maximum projection of the same trial overlaid with ROIs from CNMF spatial decomposition shows segmentation of glomeruli for a single FOV using CNMF spatial decomposition. The spatial decomposition of the FOV results in 26 glomeruli (4 dropped units after merge analysis not pictured) as outlined and numbered. c) Shows the mean traces of each glomerulus’s CNMF temporal decomposition within each flow condition (left to right: low, medium, high). Trials sorted by magnitude of normalized mean response during odor exposure. d) The deconvolved CNMF response of a single glomerulus (pink fill) to all low (grey) and high (black) flow trials across the recording sessions shows glomerular responses vary with the unique odor concentration dynamics of each plume. e) Deconvolution increases temporal accuracy of glomerular responses as shown by the mean deconvolved traces of the corresponding glomeruli depicted in (c). f) Sum of mean responses calculated within each flow condition. Mean responses (deconvolved) vary significantly between conditions (*t*(110) = 11.43, *p <* 0.001).

For each glomerulus, the average response mean (average of mean trace timepoints) was also calculated within low and high conditions. A paired samples t-test found that glomerular response means varied significantly across low and high flow conditions (*t*(110) = 9.71, *p <* 0.001). Mean responses were higher in low flow conditions, during which lower airspeed resulted in plume dynamics that were less intermittent, as shown by lower skewness and asymmetry in the deconvolved odor signals of low flow trials as compared to high flow trials ((*F* (157) = 52.7, *p < .*001 and *F* (157) = 54.9, *p < .*001 respectively)(figure 1E-H). In high flow conditions, increased intermittency produced more brief, high concentration fluctuations followed by blanks, or periods without odor signal. This decreased response mean was not due to a decrease in odor concentration means though as plumes had higher concentration means during high flow trials (*m* = 202 a.u) as compared to low flow trials(*m* = 146 a.u) (*t*(116) = 3.30, *p <* 0.001). (As the deconvolved traces are normalized using within trial averages, stimulus mean during high and low flow trials are calculated using the raw sensor signal for relative comparison.) Thus, the cumulative MT activity increased in low flow trials even as the cumulative exposure to the stimulus decreased, suggesting the response of glomerular populations are moderated by plume dynamics.

### Correlation between stimulus and glomerular activity

To determine if plume dynamics could be moderating the glomerular population response, cross-correlation was used to quantify the relation between odor concentration dynamics and simultaneously recorded glomerular activity (figure 5). Most glomeruli significantly followed plume dynamics when correlation between neural activity and odor activity was calculated across all trials (100/111), across low flow(97/111) trials, or across high flow(100/111) trials. Significant tracking of the stochastic changes in odor concentration across plume presentations is determined by comparing mean correlation coefficients to a null distribution created using a trial shuffled bootstrap analysis (see methods)(Supplemental Figure 7C). High correlation coefficients are not observed when glomerular responses are trial shuffled and ethanol recordings are no longer compared to the glomerular responses they elicited (figure 5A-B). This shows that it is not the statistics of stimulus presentations that drive this correlation, but rather the plume’s temporal dynamics unique to each trial.

**Figure 5.**
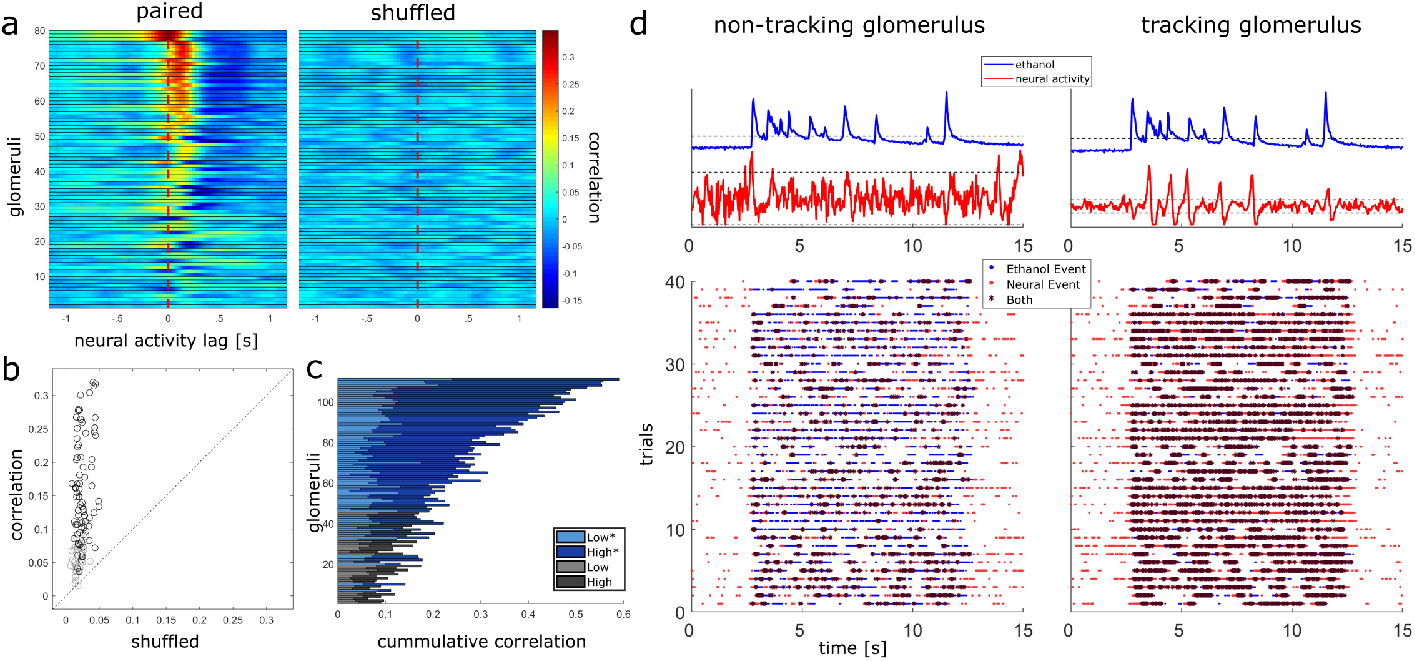
Glomerular population activity follows odor concentration dynamics across plume encounters. a) (left) The cross-correlation between the deconvolved ethanol trace and each glomerulus’s deconvolved activity trace is calculated within each trial and then averaged across trials. Each row is a glomeruli and each time point represents the cross-correlation at the indicated lag. Glomeruli are sorted in order of decreasing magnitude of correlation coefficient (see methods). (right) Same but glomerular responses are trial shuffled so that the signals compared are not from the same trial. Glomeruli are sorted to match their corresponding unshuffled cross-correlation in the right panel. b) Scatterplot of the correlation coefficient of all glomeruli compared to their respective shuffled coefficient. Glomeruli plotted in (a) are marked in black and their coefficient exceeds their shuffled coefficient from a single trial shuffled comparison by 2 standard deviations. c) Cumulative scores for each glomerulus is the sum of their correlation coefficients calculated within low and high flow conditions. The cumulative correlation plot shows variation in a glomerulus’s ability to detect changes in odor concentration dynamics varies significantly between conditions (*t*(110) = 12.81, *p <* 0.001), with most glomeruli having stronger correlation coefficients in high flow trials (*indicates glomeruli plotted in (a)). d) Binary cross-correlation. Top: Simultaneously recorded signals shown for two example glomeruli responding to the same example trial’s odor plume. Odor and glomerular activity traces plotted with their respective thresholds (dotted, odor threshold: mean during plume presentation, neural threshold: 2 st dev of baseline). Bottom: Resulting binarized traces plotted for each trial illustrate the magnitude of concurrent activity as events (stars) between the plume and the response of each glomerulus across the experimental session.

Correlation coefficients increased from the null expectations by 0.13 *±* 0.08 (mean *±* st dev) across all flow conditions, 0.08 *±* 0.07 within low flow, and 0.16 *±* 0.11 within high flow. Glomeruli were significantly better at tracking plume dynamics in high flow than they were in low (*t*(110) = 12.81, *p <* 0.001) with average correlation coefficients increasing by 0.11 *±* 0.09 (mean *±* st dev). We wondered if this increase in correlation could result from increased sparsity. Indeed correlation between two signals where the values are constant or zero for most of the time is automatically high, even if the peaks are entirely uncorrelated. If this was the case, the shuffled correlations in high flow should be significantly higher than in low flow, but this is not observed when looking at the confidence intervals for the null correlation coefficients computed within flow conditions (supplementary figure 7C). Thus, a large fraction of the glomerular population follows fluctuations during plume encounters, and the degree of dynamic tracking is moderated by plume dynamics, becoming stronger on average during plumes with higher levels of intermittency (figure 5C).

### Plume fluctuations structure glomerular network dynamics

Glomerular activity is moderated by changes in plume structure. We measured glomerular responsivity and response power to see the effect of flow condition on these measures and whether these measures were related to how well a glomerulus followed plume dynamics.

We found that there was a significant effect of flow condition on glomerular responsivity. Responsivity is defined as the proportion of trials to which a glomerulus responded to the odor (figure 7) (see methods). Glomeruli had significantly higher responsivity during low flow trials(*t* = 12.1, *p <* 0.001), with repsonsivity scores increasing by 0.21 *±* 0.18 (mean *±* st dev) as compared to high flow. Therefore, glomeruli responded to low flow trials more reliably than they responded to high flow trials. As noted previously, the average correlation between plume dynamics and MT activity increased in high flow conditions, so although glomerular responses became less reliable as airflow increased, they became more correlated with plume dynamics. Thus, plume structure changes glomerular responsiveness and response nature. Responsivity is a thresholded measure that determines if a glomerulus is more active than expected by chance during a plume presentation and does not capture the dynamics present in the response. The strength of the dynamic activity of glomeruli was determined by measuring the change in cumulative response power between baseline periods and plume presentations (see methods). Fast Fourier transform was used to measure response power within 0-5 Hz, a frequency range relative to the stimulus dynamics (figure 6). Response power (0-5 Hz) increased significantly from baseline during plume presentations (*t*(110) = 20, *p <* 0.001) by 5.7 *±* 3.0 a.u. (mean *±* st dev), a 448% increase (figure 6). On average 86.7% *±* 4.9% (mean *±* st dev) of cumulative stimulus power for each session was within 0-5Hz. The majority of cumulative response power for each glomerulus, 88.5% *±* 7.9% (mean *±* st dev), was also found to be within this range. Thus, the majority of response power for each glomerulus was measured to be within a relative response range of the stimulus (figure 6A-B). Across all glomeruli recorded,response power was not significantly different between flow conditions. To examine the effect of flow conditions on the cells that most strongly responded to the odor, we next analyzed only glomeruli whose mean response was above the 75^*th*^ percentile. Within this group of glomeruli, response power did change significantly between flow (*t*(27) = 5.52, *p <* 0.001), with stronger response power during high flow conditions. This increase reflects the significant increase in stimulus power (0-5 Hz) observed in high flow as compared to low flow trials (*t*(116) = 31, *p <* 0.001) (Supplemental Figure 6). Thus, the response power of glomeruli with the strongest signals was significantly affected by flow condition.

**Figure 6.**
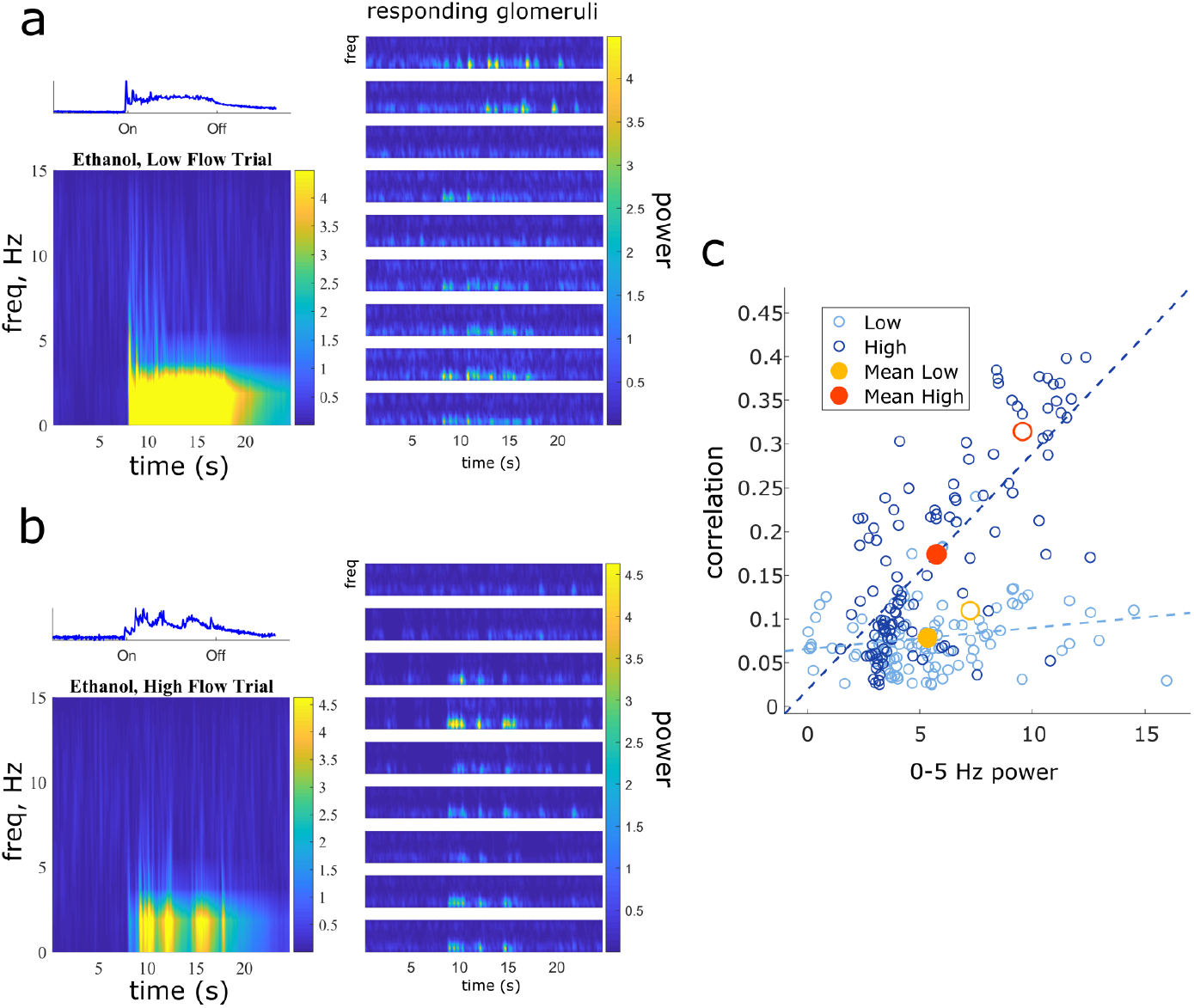
Higher magnitude of glomerular response power (0 5*Hz*) is associated with higher correlation with plume dynamics. a) Short-time Fourier transforms of a single low flow trial and a sample of responding glomeruli show most response power of the glomeruli and odor signal is concentrated between 0 5Hz. Glomeruli are sorted by increasing correlation with plume dynamics. b) Same but for a single high flow trial from another example FOV. c) With both high and low flow conditions, correlation of the glomerulus with plume dynamics is plotted against its corresponding increase in power spectrum activity between ‘odor off’ (top) and ‘odor on’ (bottom) periods. Glomeruli with higher correlation coefficients have a stronger increase in response power during plume presentations (*r* = 0.74, *p <* 0.001). When calculated within flow, this relationship is significant within high flow (*r* = 0.73, *p <* 0.001), but not within low flow(*r* = 0.19, *p* = 0.05). The average repsonse across all glomerular is plotted (red/yellow) to represent the population response. Mean response power of the glomerular population is not significantly different between low (yellow dot) and high flow (red dot), except for when calculated amount glomeruli whose mean acitvity is in the 75th percentile (low = yellow circle, high average = red circle).

There exists a relationship between each of these two response features, responsivity and response power, and how well a glomerulus follows plume dynamics. Across all trials, glomeruli with higher responsivity to plume presentations were better at following changes in odor concentration (*r* = 0.76, *p < .*001). Thus, the more reliably a glomerulus responded to plume presentations, the more likely it was to follow changes in odor concentration. This is not a perfect relationship as glomeruli that are highly responsive to the plume but not its dynamics exist (figure 7), but a glomerulus with higher responsivity is more likely to be correlated to plume dynamics than one with lower responsivity. Also, as mentioned previously, a glomerulus’s average responsivity level is moderated by flow condition. Response power was also correlated with how well the glomerulus followed changes in odor concentration, when averaged across all trials, glomeruli with higher response power were significantly better at following plume dynamics (*r* = 0.74, *p <* 0.001). When this relationship was examined within flow conditions (figure 6C), high flow was significantly correlated (*r* = 0.73, *p <* 0.001), but low flow was no longer significantly correlated (*r* = 0.19, *p* = 0.05) (All flow includes medium flow, which is not analyzed in the between flow comparisons). In addition, for a subset of strong responding glomeruli, this response power was moderated by flow condition. These results suggest that both the reliability and the temporal pattern of MT activity is significantly moderated by the odor concentration dynamics of the incoming stimuli.

**Figure 7.**
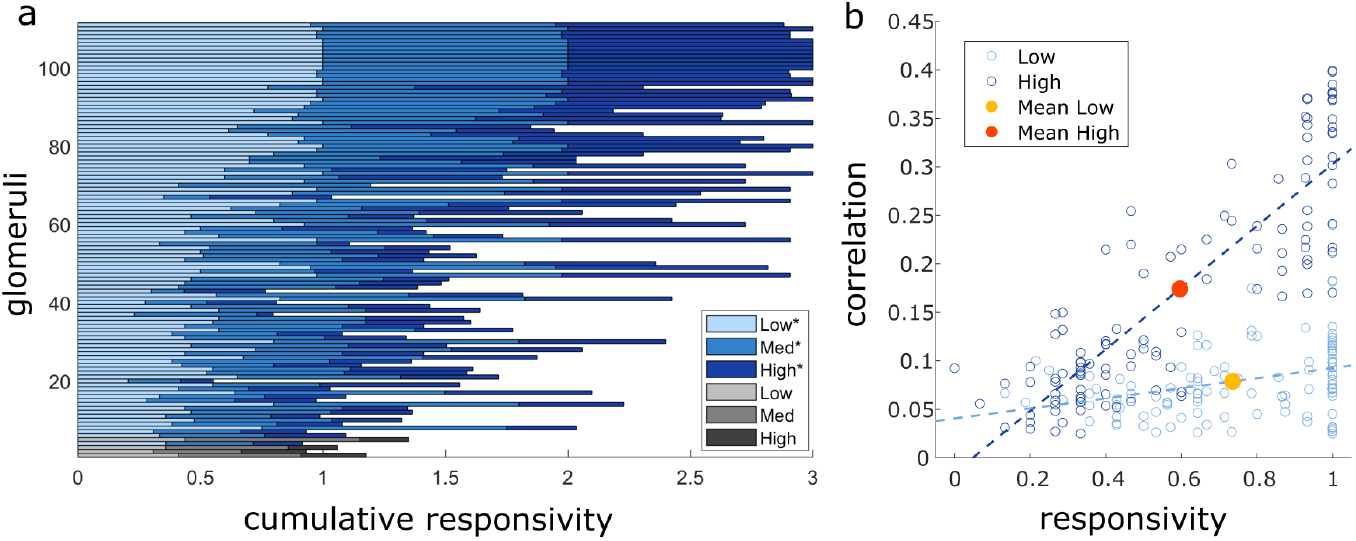
Glomeruli that respond more reliably to plumes are more correlated with their dynamics. a) The cumulative responsivity is the sum of the responsivity scores calculated within each flow condition. Glomeruli are sorted by decreasing average correlation with plume dynamics. Glomeruli whose correlation coefficient exceeds its null confidence interval (see methods) are plotted in blue hues and the remainder of the glomeruli are plotted in grey hues. This shows the magnitude of odor concentration tracking is correlated with (*r* = 0.76, *p < .*001), but not strictly defined by, response reliability as glomeruli exist that respond strongly to odor presence but not to concentration dynamics. b) Within flow condition, repsonsivity is plotted against correlation with odor dynamics for each glomerulus (circles) and for the population average across all glomeruli (dots). Across glomeruli, responsivity is positively correlated with tracking ability as illustrated by the lines of best fit. For each glomerulus, their relative responsivity decreases significantly during high flow (dark blue) as compared to low flow (light blue) (*t*(110) = 12.1263, *p <* 0.0001). To represent the population response, the average responsivity across all glomeruli is plotted against average correlation with plume dynamics (dots) within both conditions illustrating how flow moderates these relationships. For a given glomerulus, higher flow predicts a decrease in its relative responsivity level (*t*(110) = 12.1263, *p <* 0.0001) and an increase in its relative tracking ability.

These results show that the spatiotemporal dynamics of plumes play a role in structuring activity in the first olfactory relay of the mouse’s brain during natural olfactory processing.

## Discussion

Mice are adept at olfactory guided search despite the stochasticity and complexity of odor plumes used in navigation. Spatiotemporal cues present in natural odor scenes are thought to drive decision-making in olfactory search ((Mafra-Neto and Cardé, 1994); (Vickers, 2006)), (Pang et al., 2018), but how they moderate population activity in the olfactory bulb (OB) is unknown. Releasing odor within a custom-built wind tunnel, we were able to hold constant all properties of the odor stimulus and the animal’s position relative to the source and vary only the air velocity through which the plume traveled. By using this approach we altered the Reynolds number of the flow and created plumes with varying statistical structures and odor concentration dynamics. In this way, the effect of plume dynamics on MT population activity could be examined using naturally evolving odor plumes. Recording MT activity in mice expressing GCaMP6f, we show that a significant fraction of glomerular populations of MT cells follow odor plume dynamics. Additionally, the strength with which they do so is moderated by airflow, such that increased flow velocity and turbulence (Reynolds number) results in increased correlation of MT cell activity with plume dynamics. This work shows that plume dynamics structure the activity of the OB, the first relay of olfactory coding in the mouse’s brain.

The recent history of an odor stimulus has been shown to be present in olfactory encoding in both serial sampling of odor concentration in mice (Parabucki et al., 2019) and tracking of odor concentration in invertebrates (Geffen et al., 2009), showing odor concentration changes influence olfactory encoding. Although inter-sniff comparisons in mice show that MT cells can detect the sign and magnitude of changes in odor concentration (Parabucki et al., 2019), it is unknown whether they are able to resolve the dynamics of natural plumes, which span across a range of temporal scales. If odor concentration dynamics are resolved, computational work has shown that they are informative for olfactory search ((Baker et al., 2018); (Gumaste et al., 2020)). To avoid the complexity of stochastic odor plumes, the averaging of odor concentration dynamics could be an alternative strategy to navigate olfactory environments. Mean odor concentration levels are moderated both by the distance from an odor source and how close an animal is to the central stream of the plume (Crimaldi and Koseff, 2001). While this measure is potentially informative, it does not by itself sufficiently inform decision-making on the timescale observed in rodents (Gumaste et al., 2020). Therefore, it is likely that the mouse relies upon spatiotemporal features of the plume for olfactory search as information can be extracted from odor concentration dynamics (Baker et al., 2018).

Our study found a correlation between MT activity and odor concentration dynamics during plume presentations. The temporal information conveyed by MT cells could support a variety of navigation algorithms. For instance, two important dynamic features are the length of odor encounters, whiffs, and the timing between odor encounters, blanks. Whiff and blank duration are moderated by the distance between an animal and the odor source. As an animal approaches an odor source, plume encounters become shorter and more frequent ((Wright and Thomson, 2005); (Celani et al., 2014)). Blank duration has been shown to be particularly informative even when olfactory environments change. Computational modeling of olfactory search in invertebrates ((Park et al., 2016); (Rapp and Nawrot, 2020)) as well as fluid dynamics modeling (Celani et al., 2014) shows that the time between odor encounters, blank duration, is less sensitive to environmental conditions such as plume velocity or potency of the odor source that are know to affect interpretation of odor concentration dynamics ((Connor et al., 2018); (Webster and Weissburg, 2001)). Specifically, Park et al., found blank duration to be a more efficient source of information for olfactory search than instantaneous tracking of odor concentration. In our study, we observed that MT activity was more correlated with plume dynamics in high flow trials than low flow trials. Odor concentration in low flow trials was less skewed, meaning that these trials had lower intermittency and odor concentration tended to fluctuate around a central value. Alternatively, high flow trials were more skewed and were characterized by a whiff and blank structure. The fact that correlations are higher in high flow suggests MT activity may be more responsive to whiff and blank features as opposed to tracking fine fluctuations in odor concentration across more constant plume encounters. Future studies exploring the effect of a broader range of intermittency levels on MT activity during plume encounters could help determine which spatiotemporal temporal features of intermittency are moderating MT responses to plume dynamics.

The majority of glomerular activity responding to concentration dynamics could be considered to be inefficient when the OB has to perform other tasks such as odor identification and segmentation. Glomerular spatial maps, i.e. glomerular ensembles consistently responding to an odor, are thought to be one of the primary means of odor identification (Wachowiak and Shipley, 2006). Odors maps vary with concentration ((Xu et al., 2000); (Wachowiak and Cohen, 2001)), but are stable enough to reliably encode odor identity (Belluscio and Katz, 2001). Although pulsed odors can be rapidly discriminated (*<*200ms, or a single sniff, (Uchida and Mainen, 2003)), in a natural olfactory environment where odors are intermittent ((Celani et al., 2014); (Murlis et al., 2000)) and mixed, identification becomes a much more complicated task especially for identification of mixtures. Glomerular ensembles reliably responding across odor encounters could aid odor discrimination in mixed odor environments. Spatial maps of odor identification will overlap in natural olfactory scenes where an animal encounters signals from multiple odor sources as it navigates a plume. Odors co-released travel together (Celani et al., 2014), and therefore a mixture of odors emanating from the same source will have correlated temporal dynamics in the plume and will thus be experienced by a searcher as having correlated encounters across whiffs. This means the probability of the signal from separate sources arriving together reliably across whiffs would be low if odors are released from spatially separated sources. Grouping and demixing these odor representations using the correlation, or lack thereof, in the odor concentration dynamics could aid odor discrimination in complex environments (Hopfield, 1991).

Since our studies are not recorded at the individual cell level, the potential degree of heterogeneous tuning to different features among MT cells within a single glomerulus was not examined. It could be that observed correlations of glomerular MT populations were a product of the collective activity of heterogeneously tuned MT cells within a glomerulus, but MT responses have been shown to linearly sum odor inputs (Gupta et al., 2015), which contradict the idea that they are directly tuned to different features of plume dynamics. At the same time, this does not infer MT activity responding to plume dynamics is homogeneous as the responsiveness (Adam et al., 2014) and the response (Geramita and Urban, 2017) varies between MT cells across concentration levels (Cleland and Borthakur, 2020). Future research across a variety of odor concentration dynamic regimes and odor mixtures at both the cellular and population level are needed to further investigate the degree to which bulbar responses are tuned to features of odor concentration dynamics and how this tuning may impact optimal encoding of odor information.

Our data show that MT activity in the OB of mice follows the temporal dynamics of odor plumes. Additionally, we demonstrate that this effect is stronger under conditions that generate larger Reynolds numbers. Following odor concentration dynamics within plumes could enable MT cells to convey information useful for olfactory search. Following the temporal dynamics of odor plumes may also be an efficient form of multiplexing odor identity and source location for the first olfactory relay in mice. Although MT activity responds to changes in odor concentration, the observed correlations do not suggest perfect tracking at the level of individual glomeruli and indicate inter-individual differences in the degree to which glomeruli follow plume dynamics. Future research focusing on location encoding across a wide range of both intermittency regimes and odor panels is needed to clarify the degree to which bulbar activity is tuned to features of plume dynamics and how a balance between identity coding and concentration coding is instrumental in supporting the wide variety of behaviors enabled by olfaction.

## Supporting Information

**Figure S1.**
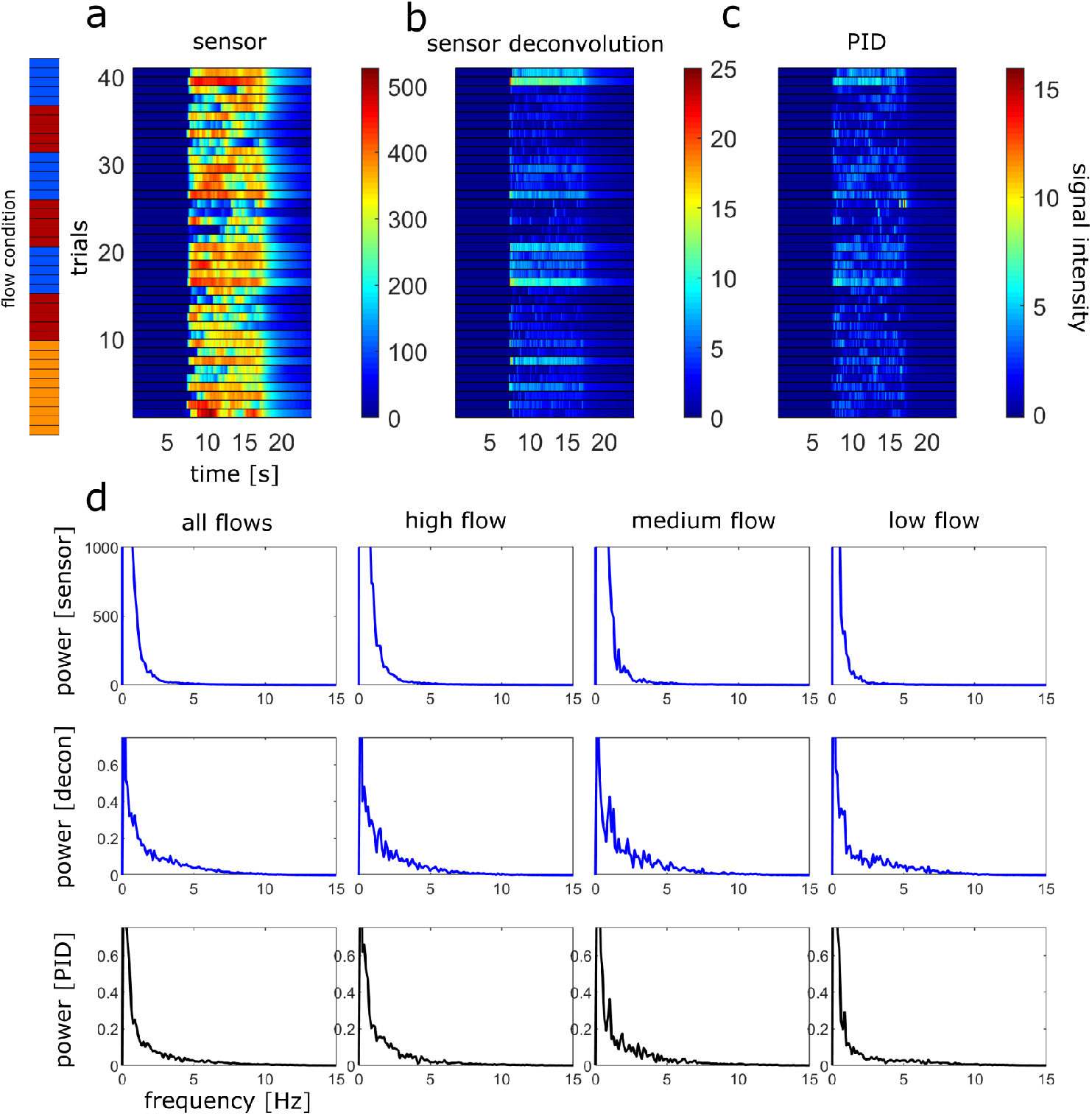
Deconvolved Ethanol signal and spectral decomposition from one session divided by flow type. a) The raw ethanol sensor output is shown for all trials within a single session. (red, orange, and blue are high, medium and low flow respectively). b) The deconvolution of the raw ethanol signal. c) The simultaneously recorded signal from a co-localized PID. Sensor deconvolution and PID are significantly correlated as calculated during plume presentations (*r* = .3089, *p < .*001). d) (top) The spectral decomposition calculated across all trials or within flow conditions for the raw sensor signal shown in (a). Only the middle 8 seconds of the 10 second plume presentation are analyzed to avoid onset or offset dynamics. Spectral decompositions are also plotted for the ethanol deconvolution (middle) shown in (b) and for the PID (bottom) shown in (c).

**Figure S2.**
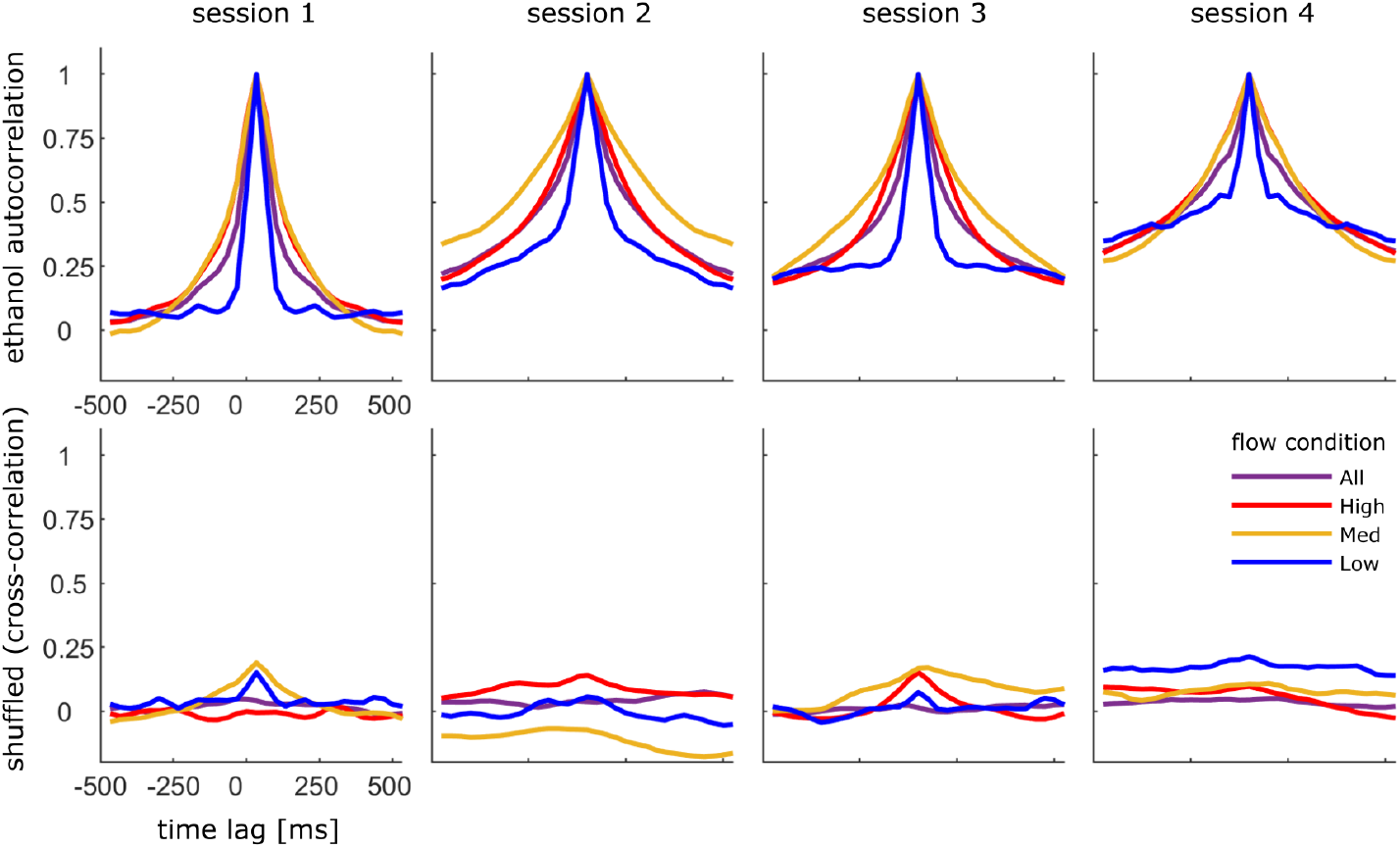
Normalized autocorrelation of ethanol trace. a) The Normalized average autocorrelation of the deconvolved ethanol signal is shown for each of the 4 experimental sessions. Traces are calculated across all trials within the designated flow conditions. b) Same as (a) but trials are shuffled such that each cross-correlation is calculated between the deconvolved ethanol signal from two different trials.

**Figure S3.**
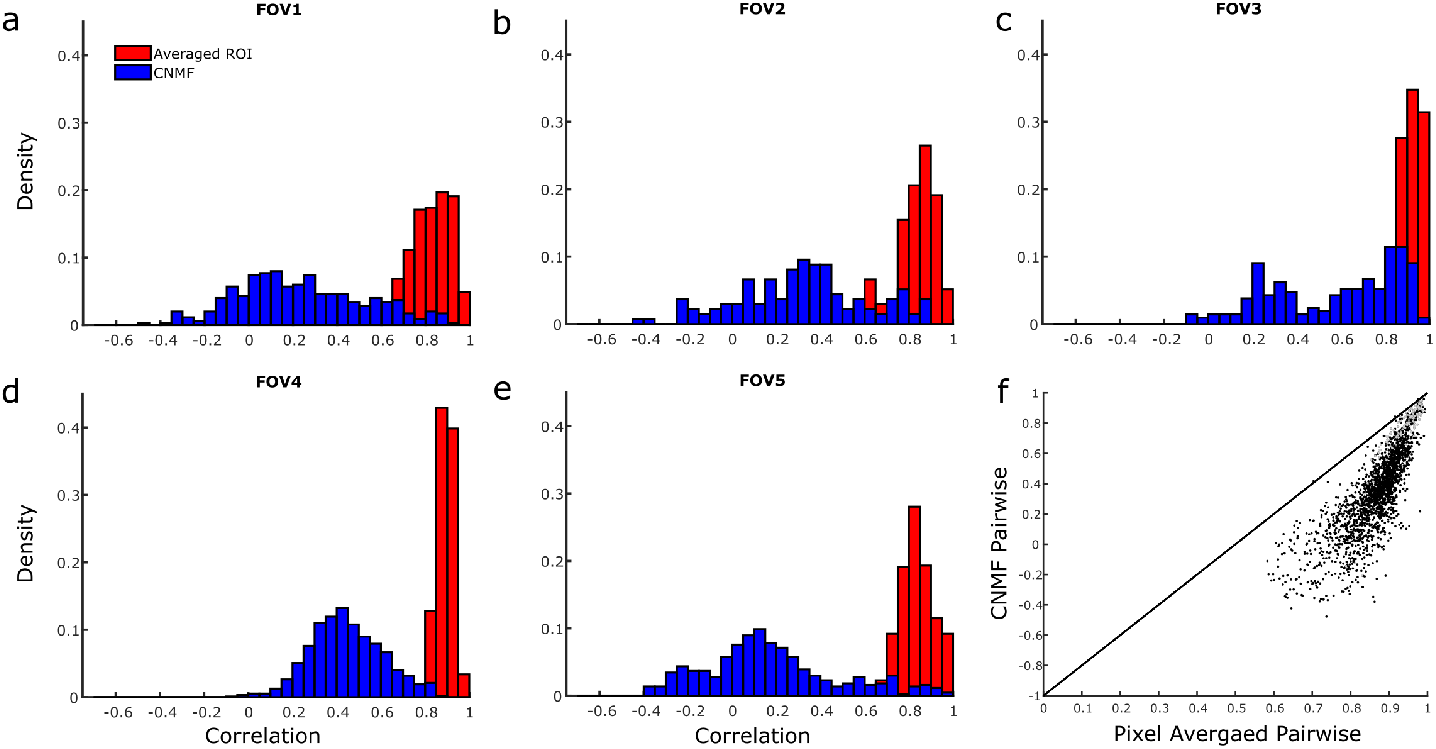
Pairwise correlations between glomerular ROIs are decorrelated by CNMF. a-e) Distributions of the pairwise linear correlation coefficients from all possible glomerular pairs within each FOV. Correlations are calculated on the temporal decomposition of CNMF ROI signals (without deconvolution) which include denoising and demixing (blue) or on the raw pixel values as averaged within the same ROI boundaries (red). Pairwise correlations are calculated between the two glomeruli during stimulus presentation. f) Scatterplot of each pairwise correlation shows CNMF de-noising decorrelates the signal for each pair of glomeruli. Pairwise correlations in grey are from ROI pairs that have been considered to originate from the same glomerulus after the merging threshold analysis (Supplemental Figure 4).

**Figure S4.**
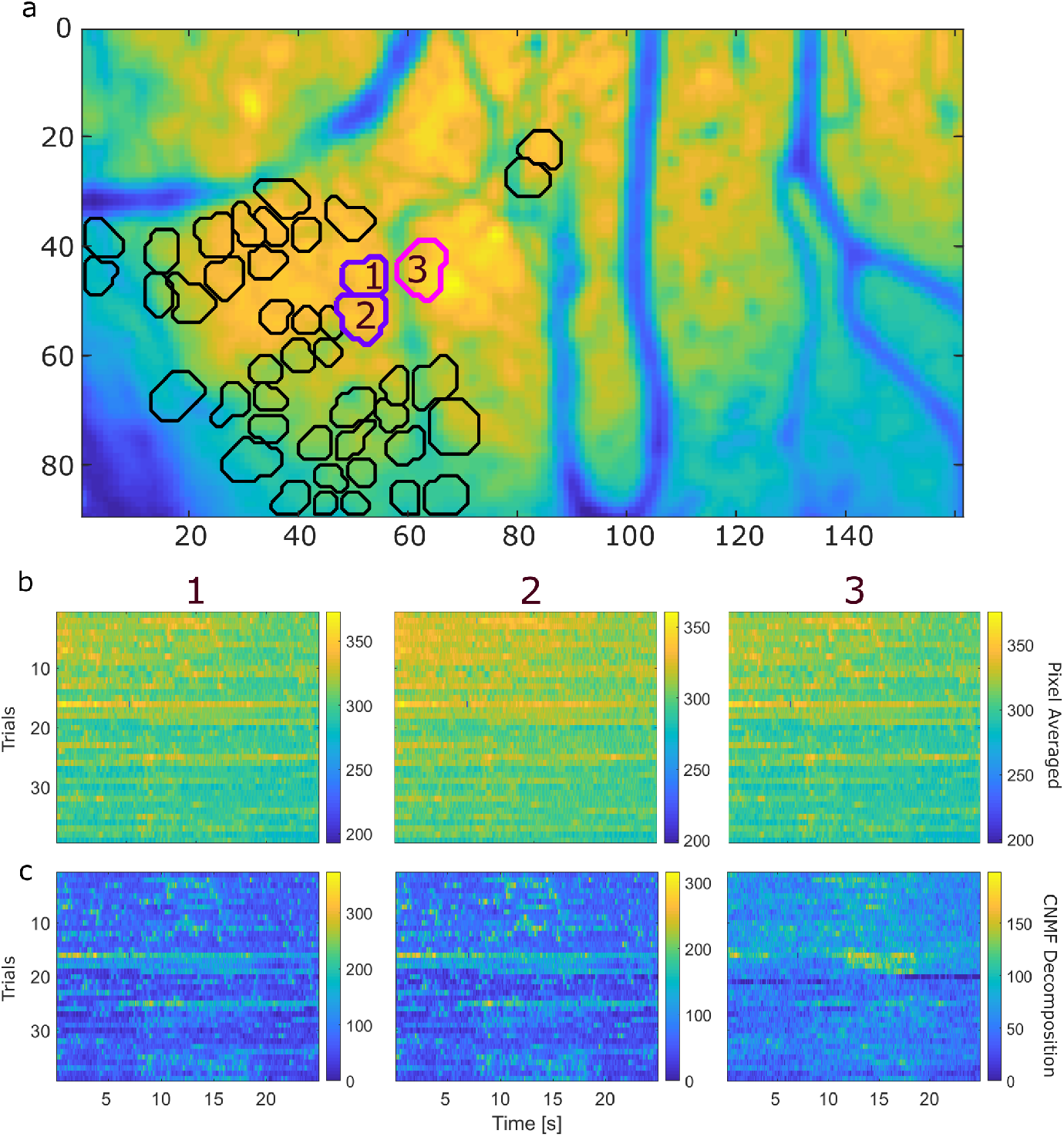
Detecting oversegmentation of ROIs after CNMF decomposition. a) ROI segmentation of glomeruli for FOV 4 using CNMF spatial decomposition of activity across an entire session. ROIs circled in purple do not meet thresholds for independent ROIs (methods) and thus are considered to originate from the same glomerulus, while the ROI in magenta is considered to be a separate glomerulus. b) Raw fluorescent signal averaged across all pixels within each of the 3 ROIs (without denoising or de-mixing) is plotted for all trials in the session. Numbers above each plot refer to ROI numbering in panel (a). c) CNMF signal across all trials plotted for the corresponding ROIs in (b).

**Figure S5.**
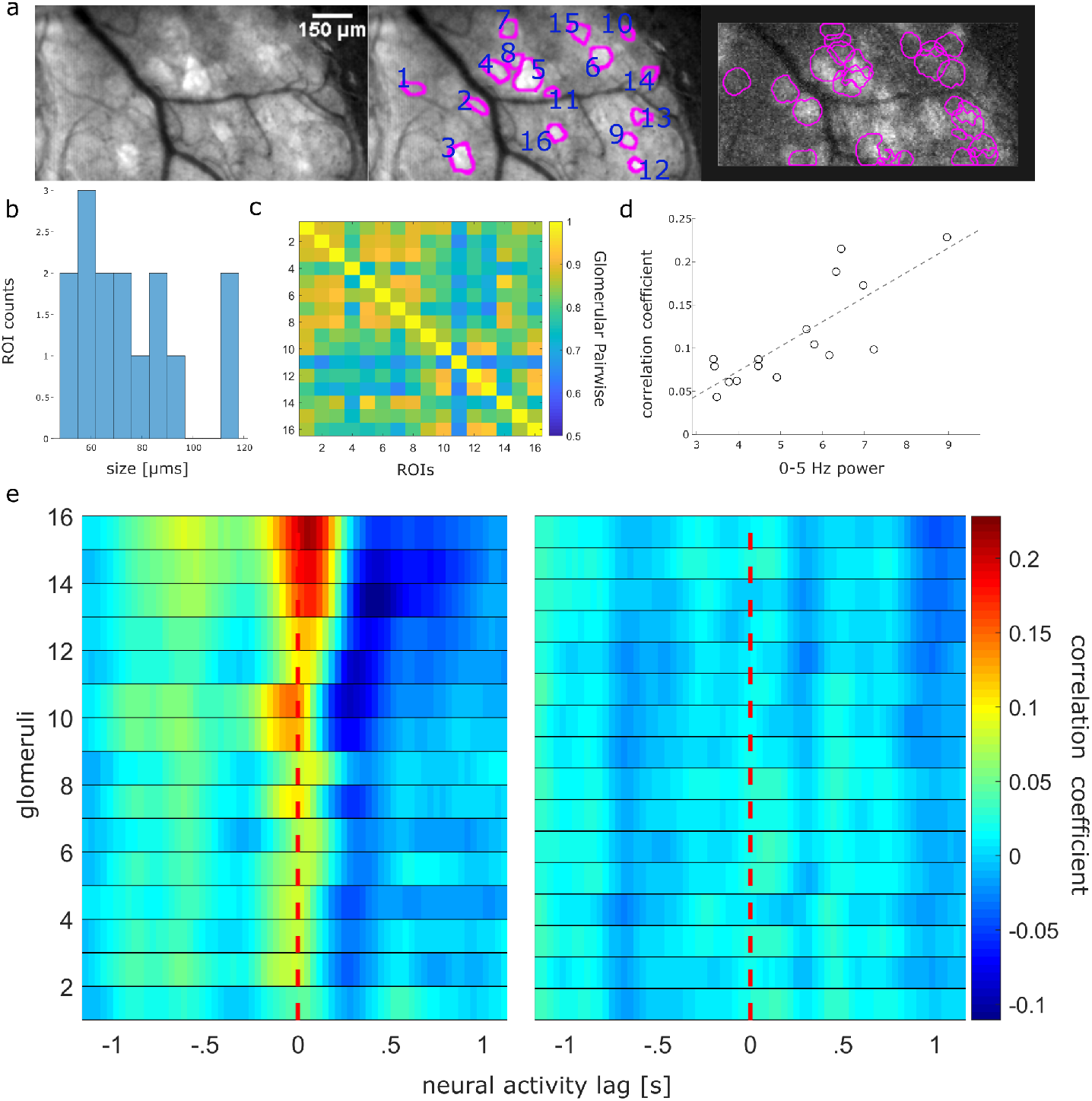
Hand-drawn ROIs have qualitatively similar results but higher pairwise correlations. a) (left) St dev projection of a single FOV during the first trial of the session. (middle) ROIs from hand-drawn analysis are plotted over the projected st dev (left). ROIs determined using recordings and st dev projections averaged across trials within each flow condition. (right) Mean subtracted maximum projection of the same trial overlaid with ROIs from CNMF spatial decomposition. b) Distribution of hand-drawn ROI sizes from (a). c) Trial averaged linear pairwise correlation coefficients of ROI activity during plume presentations shows high correlations between ROIs. d) Glomerular correlation with odor dynamics is plotted against their corresponding change in repsonse power (0 5Hz) between ‘odor off’ (d) and ‘odor on’ periods showing glomeruli with higher correlation coefficients have higher activity power (*r* = 0.80, *p* =*<* 0.001). The dotted line plots the line of best fit using OLS regression. e) Glomerular activity is correlated with stimulus dynamics at varying lags (right) with all mean coefficients exceeding a 95% confidence interval of their shuffled mean coefficients. Each row is a glomerulus and each time point represents the mean correlation coefficient between odor and response deconvolutions at the indicated lag. Glomeruli are sorted in order of decreasing correlation. (right) Same but neural activity responses are shuffled so that the signals compared are not from the same trial. The glomeruli are sorted to match their corresponding unshuffled cross-correlation in the right panel.

**Figure S6.**
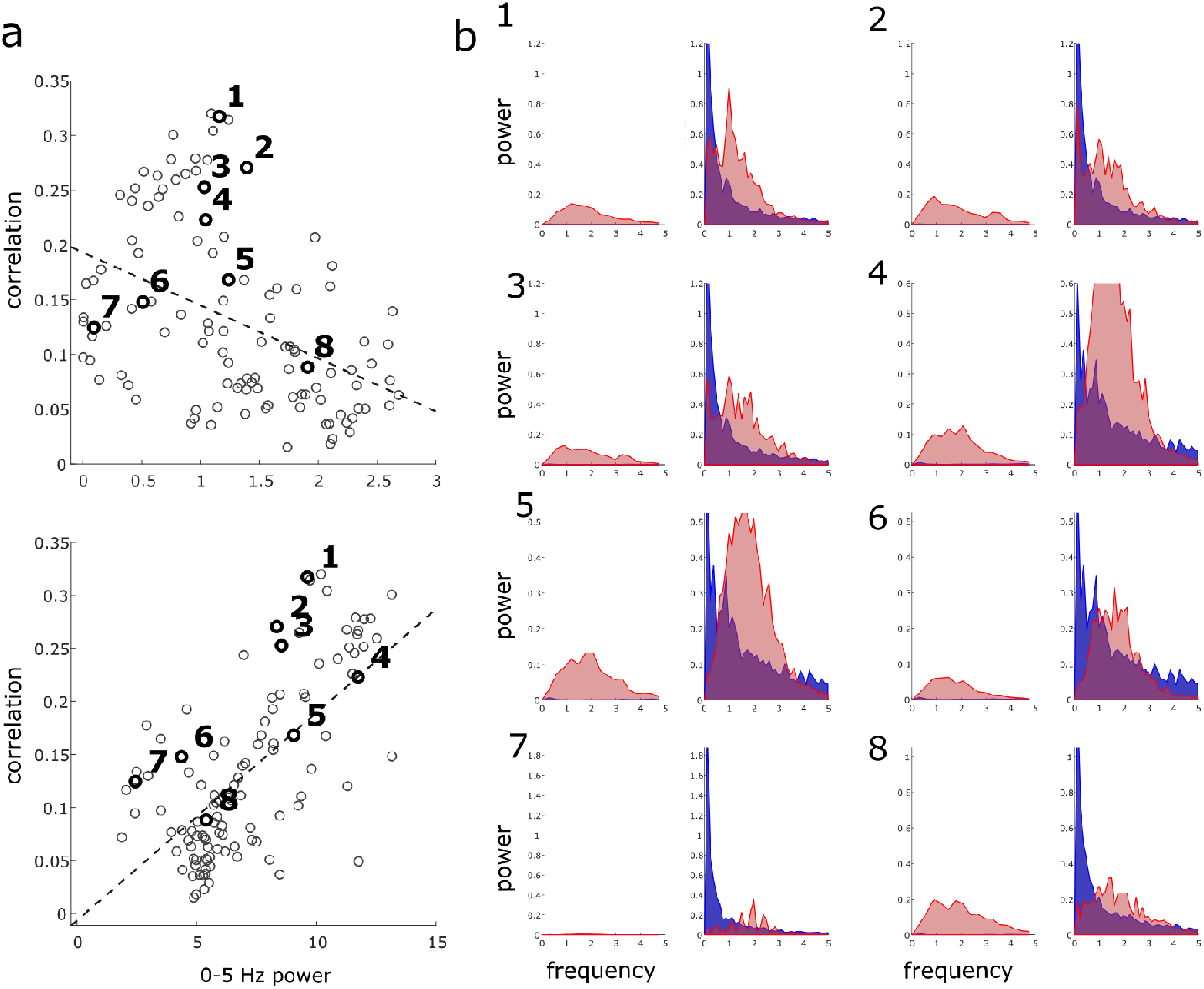
0-5 Hz Power spectrums for odor and glomerular responses. a) Average binary cross-correlation (subplots same as shown in Fig. 6) for each glomerulus is plotted against the power spectrum of its activity during ‘odor off’ (top) and ‘odor on’ (bottom) periods. b) The power spectrums of the deconvolved odor (blue) and glomerular (red) traces are plotted for 8 example glomeruli (numbered accordingly in (a)) during ‘odor off’ and ‘odor on’ periods.

**Figure S7.**
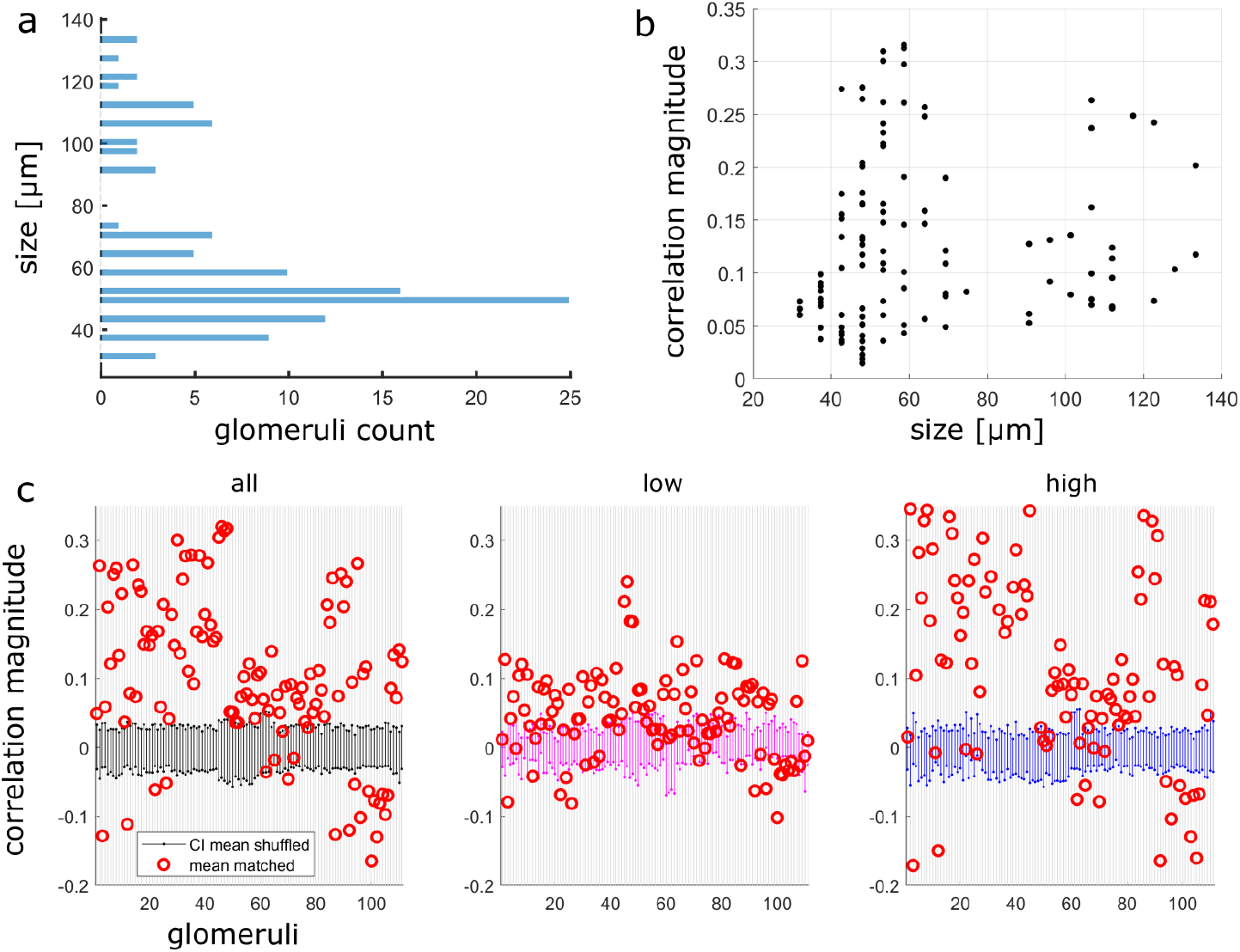
Glomerular sizing and plume following behavior. a) Distribution of CNMF ROI sizes for all sessions (*μm*). b) ROI size (*μm*) plotted against the mean correlation coefficient between deconvolved glomerular and ethanol signals during odor presentation shows correlations between ROIs and plume dynamics are not driven exclusively by singular MT cell activity. Kendall’s tau coefficient between size and odor concentration tracking (*r*_*τ*_ = 0.1563, *p* = 0.202) shows that correlation to plume dynamics is not exclusive to or related to smaller sized ROIs (ones similarly sized to individual MT cells). This suggests response to plume dynamics across plume encounters is also occurring at the glomerular level. c) Mean correlation coefficients of glomeruli (red) are plotted against their respective bootstrapped 95% confidence interval of null mean coefficients (see methods for trial shuffled bootstrap analysis). Null confidence intervals are calculated across all flow conditions (black), within low (pink) or within high (blue) flow.

## Author Contributions

Author contributions: DHG, AG, and SML initiated the study and designed the experiments. DHG and SML built the head-fixed imaging setup, built the plume delivery setup, and wrote acquisition software. SML and LX performed cranial window surgeries. SML and LX performed experiments. DHG, MS, and SML performed analysis of neural data. DHG, AG, SML, and NR performed analysis of plume dynamics. DHG, AG, NR, and SML developed deconvolution based on Tariq et al., 2019 (MT) and MT assisted with sensor characterization. DHG, AG, SML and NR generated figures reporting data with input from MS. All authors helped with the text.

## Acknowledgments

We would like to thank Michael Grybko for in vitro characterization of GCaMP6f expression in MT cells, Katie Ferguson (Postdoctoral Fellow, Neuroscience, Yale University) for consultation regarding calcium imaging data acquisition and analysis, Kelly Chang (Graduate Student, Psychology, University of Washington) for advice on data analysis and figure design, and Jin Young Park for assisting in data collection.

